# Spectral consistency in sound sequence affects perceptual accuracy in discriminating subdivided rhythmic patterns

**DOI:** 10.1101/2023.05.15.540754

**Authors:** Jun Nitta, Sotaro Kondoh, Kazuo Okanoya, Ryosuke O. Tachibana

## Abstract

Musical compositions are distinguished by their unique rhythmic patterns, determined by subtle differences in how regular beats are subdivided. Precise perception of these subdivisions is essential for discerning nuances in rhythmic patterns. While musical rhythm typically comprises sound elements with a variety of timbres or spectral cues, the impact of such spectral variations on the perception of rhythmic patterns remains unclear. Here, we show that consistency in spectral cues affects perceptual accuracy in discriminating subdivided rhythmic patterns. We conducted online experiments using rhythmic sound sequences consisting of band-passed noise bursts to measure discrimination accuracy. Participants were asked to discriminate between a swing-like rhythm sequence, characterized by a 2:1 interval ratio, and its more or less exaggerated version. This task was also performed under two additional rhythm conditions: inversed-swing rhythm (1:2 ratio) and regular subdivision (1:1 ratio). The center frequency of the band noises was either held constant or alternated between two values. Our results revealed a significant decrease in discrimination accuracy when the center frequency was alternated, irrespective of the rhythm ratio condition. This suggests that rhythm perception is not only shaped by temporal structure but also affected by spectral properties.

## Introduction

We detect temporal regularity and capture complex structures of sound events in almost all styles of music. Each perceived regular pulse is called a beat. Subdividing each beat interval provides various characteristics in the rhythmic pattern of the music. In other words, we perceptually organize various sound events into a rhythm structure according to their temporal properties by detecting and subdividing regular patterns in sound sequences [1]. For example, subdividing beats with simple integer ratios such as 1:1 and 2:1 is far more frequently observed in Western music [2]. In contrast, complex ratios such as 5:3 are very common in drum ensemble music in West Africa [3]. One of the famous subdivisions is swing. Swing is a long-short duration alternating pattern considered an essential factor in jazz music. The integer ratio of swing is generally considered to be 2:1, but it approaches 1:1 as the tempo increases. [4]. This study focuses on the perceptual accuracy of such subdivision rhythms.

Various temporal factors influence our rhythm perception. We can easily perceive regular beats even though the actual timings of the performed beats usually fluctuate and deviate from an exact regular pattern [5]. Despite such variations in beat durations, audiences perceive temporal regularity in the performed sound pattern and interpret the variation as intentional aesthetic expressions [6,7]. When beats are subdivided into finer and/or uneven intervals, rhythm perception becomes more variable. Equal subdivisions cause the tempo to be perceived as slower, even while the actual tempo remains constant [8]. Conversely, sound sequences with an identical subdivision pattern but different tempi tend to be perceived as different rhythmic patterns [9]. Precise subdivision perception relies on detecting subtle relative differences among time onsets in a regular beat sequence. We can notice approximately 2.5 % displacement of one note onset in an isochronous sequence with a tempo of 240-ms inter-onset interval [10], although such detection threshold increases when the tempo becomes faster [11]. Moreover, asymmetries have been reported in which subdivisions are perceived as equally spaced, even though they are short-long patterns, and this does not happen in long-short patterns [12].

It is known that not only the temporal property but also the sound timbre, or spectral property, influences how we organize the sound sequence structure. This issue has been intensively investigated in the research field of auditory stream segregation [13,14]. For example, a study using band noise suggested that differences in the center frequency of the band noise best predicted the degree to which stream segregation occurred [15]. Another study used a sound sequence alternating between pure tone and narrow-band noise regularly and delaying only even-numbered sounds gradually [16]. The result showed that the greater the difference in spectral aspect between the two sound elements, the less likely the subjects were to notice the delay of even-numbered sounds. The authors claimed that differences in timbre enhanced the stream segregation, thereby making it more difficult to perceive the temporal relationship of sound elements across different streams. This raises the possibility that spectral variations might influence the accuracy of rhythm perception. In real music pieces, rhythmic patterns typically consist of sounds with different timbres, such as those produced by a drum set. Consequently, an experimental assessment of how varying spectral cues affects the accuracy of rhythmic pattern perception offers novel insights for understanding rhythm perception in practical musical contexts.

In the present study, we examined the effect of different timbres on rhythm perception by psychophysically measuring the discrimination accuracy of multiple rhythmic patterns, including swing rhythms. We used a sound sequence consisting of band noise bursts, and manipulated the spectral cue by changing their center frequency. Experiments were performed online via a web browser to recruit many participants. We conducted three experiments corresponding to three rhythmic patterns: long-short rhythm (Exp. 1), short-long rhythm (Exp. 2), and straight rhythm (Exp. 3). To confirm the reliability of online experiments, particularly in the aspect of data reproducibility across arbitrary sound listening environments, we additionally conducted an experiment with participants who performed their tasks in the local laboratory so that we could confirm the listening environment (Exp. 1’).

## Methods

### Participants

We initially recruited a hundred participants for each of three online experiments via a crowd-sourcing service (CrowdWorks, Inc., Japan). None of them reported a history of hearing problems. A subset of participants was excluded from the analysis (5, 6, and 14 participants in Exps. 1, 2, and 3, respectively) due to incomplete data retrieval caused by malfunctions in the online experiment system. Then, we screened the dataset according to correct rates in screening trials and goodness of fit to the psychometric function (see Analysis section). After the screening, data from 40 (20 males, 20 females; age: M ± SD = 39.2 ± 8.10 yo), 43 (21 males, 21 females, 1 other; age: 36.3 ± 8.66), and 30 (18 males, 12 females, age: 38.9 ± 8.96) participants for Exps. 1, 2, and 3 were included in further analysis (**Table 1**). We allowed participants to join multiple experiments and additionally assessed intra-individual factors across different experiments. Several people participated in two experiments: after screening, 12, 10, and 12 participants overlapped in Exps. 1-2, 2-3, and 3-1, respectively. Three participated in all experiments. We additionally conducted the Exp. 1’ on 16 people (8 males, 8 females; age: 27.5 ± 6.74) in our local laboratory without using the crowd-sourcing service so that we were able to confirm the acoustic environment of the participants.

**Table 1.** The number of participant information.

|  | Exp. 1 | Exp. 2 | Exp. 3 | Exp. 1' |
| --- | --- | --- | --- | --- |
| screened (recruited) | 40 (95) | 43 (94) | 30 (86) | 16 (16) |
| male | 20 (53) | 21 (51) | 18 (50) | 8 (8) |
| female | 20 (42) | 21 (42) | 12 (36) | 8 (8) |
| other | 0 (0) | 1 (1) | 0 (0) | 0 (0) |

We confirmed music expertise with a questionnaire. None of the participants was an expert musician, but almost half had experienced intensive music instrument practices as amateurs: 46, 47, and 46 participants in Exps. 1, 2, and 3, respectively. Note that the intensive music experience refers to training at least two hours per day and may include experience in music classes and club activities but excludes experience in compulsory music classes. All participants were given informed consent prior to the experiment. All experimental procedures were approved by Ethics Review Committee on Experimental Research with Human Subjects of the Graduate School of Arts and Sciences, The University of Tokyo (No. 718-3).

### Sound stimuli

We employed sound sequences of brief band-passed noises as stimuli (**Figure 1**). The center frequency of the band noise was manipulated to form two conditions: the same-frequency condition, in which the center frequency was fixed at 1500 Hz, and the different-frequency condition, where the center frequency was alternated between 1000 and 2250 Hz (14 semitones apart). The bandwidth was fixed at 1000 Hz. Each band noise was created from white noise by band-pass filtering (FIR, for 1000-Hz noise: 582 taps, 1500-Hz: 1020 taps, 2250-Hz: 2038 taps), which was designed by the ‘band-pass’ function in MATLAB. The duration of each noise burst was 30 ms with a rise and fall time of 10 ms.

**Figure 1.**
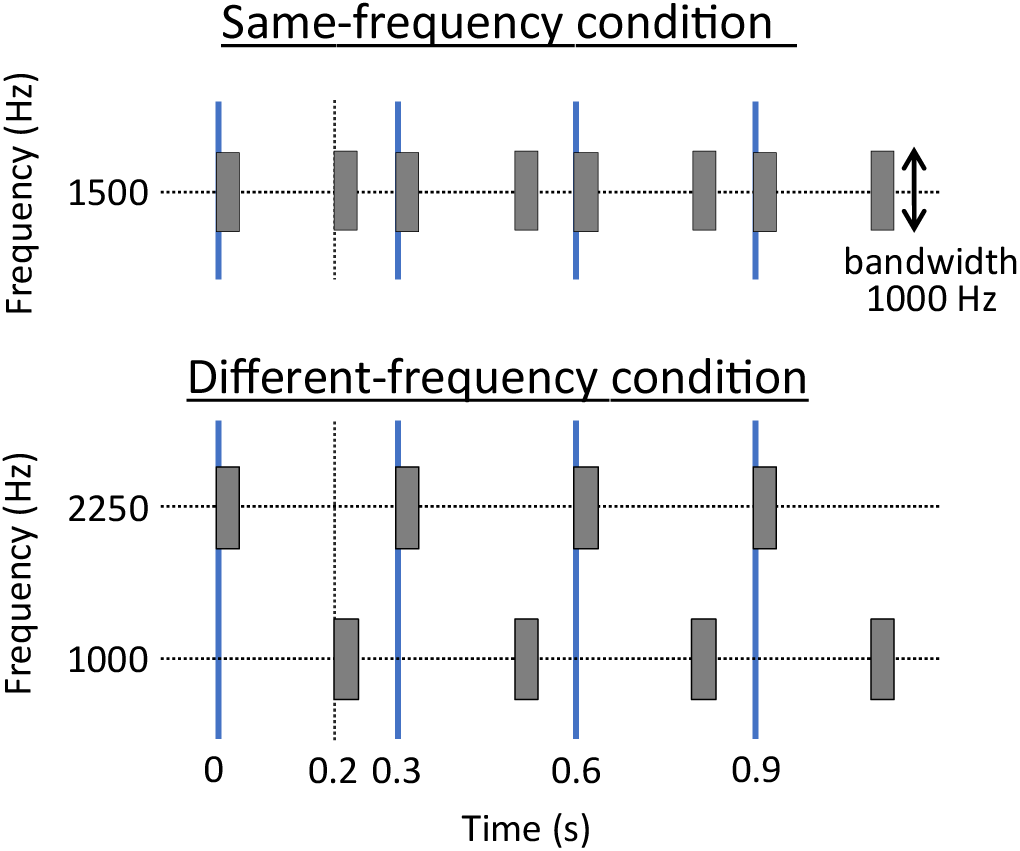
Schematic drawing of stimuli design for interval ratio *r* = 0.5. Upper indicates a stimulus used in the same-frequency condition. Lower shows a stimulus used in the different-frequency condition. The thick solid line (blue) indicates beats.

Each stimulus sequence consisted of eight noise bursts. Inter-onset intervals of odd number-th noises were fixed at 300 ms, which formed musical beats (hence, we call this inter-beat interval: IBI). Onsets of even number-th noises were varied according to an interval ratio index *r*, defined as follows:

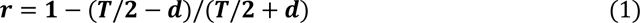

here, *T* represents IBI, and *d* shows onset displacement from the middle timepoint between beats (*T*/2). The *r* value becomes 0, 0.5, or −0.5 for the interval ratio of 1:1 (straight rhythm), 2:1 (swing), or 1:2 (inverse-swing), respectively. In the different-frequency condition, either the odd-number-th (on-the-beat) noises or the even-number-th (off-the-beat) ones had the higher frequency band (2250-Hz centered), and the others were placed at the lower band (1000-Hz centered). All sound stimuli were digitally generated at 44.1 kHz, 16 bits sampling. Their amplitudes were adjusted to have the same root-mean-square level.

The stimuli in the different-frequency condition had two different types: high-low (on-beat noises had higher center frequency) and low-high (on-beat noises had lower), though this factor was out of our interest in this study. We randomly assigned participants to the one of two conditions (each participant experienced only either high-low or low-high stimuli).

### Task and condition

In each experimental trial, two stimulus sequences were successively presented with a 300-ms interval between them. The former sequence was the standard stimulus, and the latter was the comparison. We conducted three experiments (Exps. 1, 2, and 3) in which the standard stimuli had the interval ratio of *r* = 0.5, −0.5, and 0 (corresponding to 2:1, 1:2, and 1:1), respectively. The comparison stimuli were prepared with varying *r* values in five small steps: 0.2, 0.4, 0.5, 0.6, 0.8 (Exp. 1); −0.2, −0.4, −0.5, −0.6, −0.8 (Exp. 2); − 0.3, −0.15, 0, 0.15, 0.3 (Exp. 3). Participants were asked to choose “bouncier one” (corresponding to larger absolute *r* value) from the two stimulus sequences in Exps. 1 and 2, while asked to select “equally spaced one” in Exp. 3. We randomized the order of the standard and comparison stimuli in Exp. 3 to avoid selection bias since the standard was always “equally spaced” in this experiment.

Participants were required to respond by mouse click within 2.5 s after the offset of the second stimulus sequence. After participants responded, the correctness of the answer was feedbacked as symbols ○ (correct) or × (incorrect) on the computer screen. To confirm whether participants appropriately focused on the task, we inserted screening trials in experimental sessions. In the screening trial, the interval ratio of comparison stimuli was *r* = 0.2, −0.2, or 0.3 in Exps. 1, 2, or 3, respectively, so that participants could easily choose the correct response if they kept paying attention to the presented stimuli.

### Procedure

Each of Exps. 1-3 consisted of 240 trials (2 frequency conditions × 5 interval ratio conditions × 24 trial repetitions) that were equally separated into six blocks with inserting brief rests. In Exp. 1’, the number of repetitions was increased to 40 trials (total 400 trials) to obtain stable data. The frequency conditions and onset displacement conditions were randomized within the block. Screening trials were added at the beginning and end of the first and second blocks, respectively. Before starting the experiment, participants underwent a practice session that consisted of 8 trials, one for each of the eight conditions, except for the one with an onset displacement of 0 ms. Correct/incorrect feedback was provided after each response during the practice session. When the response time exceeded the required time (2.5 s), the participant was immediately informed that the time was up.

Before the experiment session, we conducted a headphone-screening task to confirm whether the participants wore headphones. Participants were asked to select the sound with the lowest loudness among the three different sound stimuli, according to a previous report [17]. The three sound stimuli were (i) a diotic presentation of 200-Hz pure tone, (ii) a 6-dB lower 200-Hz pure tone, and (iii) a dichotic presentation of 200-Hz pure tone to one ear side and its antiphase waveform to the other side. After the experiment, we asked participants about their music experience using a questionnaire. This questionnaire included questions about musical instrument experience, daily music listening, favorite genres and songs, and dance or rhythm game experience.

All experiments were programmed using a JavaScript-based experiment builder (lab.js [18]) and conducted online via Open Lab (Open Lab Online UG, Konstanz, Germany). The entire experiment took around 30 minutes.

### Analysis

To assess whether discrimination accuracy varies with spectral cue, we classified data into the same- and different-frequency conditions, and approximated by the curves described below, respectively. For each participant, we calculated the mean response rate for each interval ratio. In Exps. 1 and 2, the obtained response rate was approximated by a sigmoid function (cumulative normal distribution). The shape of sigmoid curve is determined by two parameters: *μ* gives the point of maximal gradient, which corresponds to the point of subjective equality (PSE); and *σ* shows the steepness of the gradient, which is equivalent to the discrimination accuracy. We calculated *μ* and *σ* for each subject and compared them with frequency conditions. For Exp. 3, we fitted a simple Gaussian function with fixing *μ* at 0 and the maximum value at 0.5 instead of using the sigmoid function because the response rate in Exp. 3 were supposed to be the highest (0.5) at the standard stimulus interval ratio (*r* = 0). Additionally, we performed partial fittings for data either in positive or negative interval ratio values (0 ≤ r < 1 or −1 < r ≤ 0) to assess the positive-negative asymmetry of discrimination accuracy.

Data with one or more incorrect answers in the screening trials were excluded as participants did not respond appropriately. We set further screening criteria to maintain the data reliability as follows: (1) the adjusted coefficient of determination (*R*^2^) in the sigmoid fitting must be 0.5 or larger; (2) the absolute value of PSE (|*μ*|) must range between 0 and 1; (3) the discrimination accuracy (*σ*) must be 0 or larger. We excluded participants from further analyses if the data did not satisfy these criteria either in the same-frequency or different-frequency condition.

We examined homogeneities of variance in *μ* and *σ* data by the two-sample F test, which failed to guarantee the homogeneity in *μ* in Exp. 1 (*F* (39, 39) = 2.32, *p* = 0.010), *σ* in Exp. 1 (*F* (39, 39) = 7.88, *p* < 0.001), *σ* in Exp. 2 (*F* (42, 42) = 3.02, *p* < 0.001), and *σ* in Exp. 3 (negative: *F* (17, 17) = 7.47, *p* < 0.001; positive: *F* (17, 17) = 3.22, *p* = 0.021). Thus, we employed Wilcoxon signed-rank tests for this analysis. The significance level *α* was set at 0.05. Post hoc analysis confirmed that the statistical power (1−*β*) was more than 0.8 in Exps. 1, 2, and 3 (Exp. 1: 1–*β* = 0.99; Exp. 2: 1–*β* = 0.98; Exp. 3: 1–*β* = 0.99). Note that there were some data (*n* = 3) in which *σ* was very close to zero in Exp. 3, and we conducted the test in each case when they were excluded and not excluded.

We performed a supplementary analysis to check if the effect of high-low order, which is out of interest in the present study, was ignorable. The analysis with the linear mixed model did not find any significant difference between the high-low and low-high groups consistently across three experiments (see Support Information). Since the main focus of this experiment was the effect of center frequency on perceptual accuracy, we decided to pool data from the high-low and low-high groups.

## Result

### Reliability of the online experiment

First, we confirmed the reliability of the online experiments regarding hearing environments by comparing the results of Exp. 1 with its offline version Exp. 1’. The critical but uncontrollable factor in online experiments is the potential difference in acoustic conditions, such as the frequency characteristics of headphones and background noise from the environment. In our case, we used band-passed noise to reduce the influence of variability in the frequency characteristics of the participants’ headphones. However, it is almost impossible to control the background noise factor. Therefore, we here performed the same experiment as in Exp. 1 in the local laboratory for a subset of participants, that is, under identical background conditions. Sigmoidal approximations for Exps. 1 and 1’ showed almost exactly overlapping curves for the same-frequency condition (**Figure 2a**), demonstrating that the potential variability in listening environments did not affect the main results of our online experiments.

**Figure 2.**
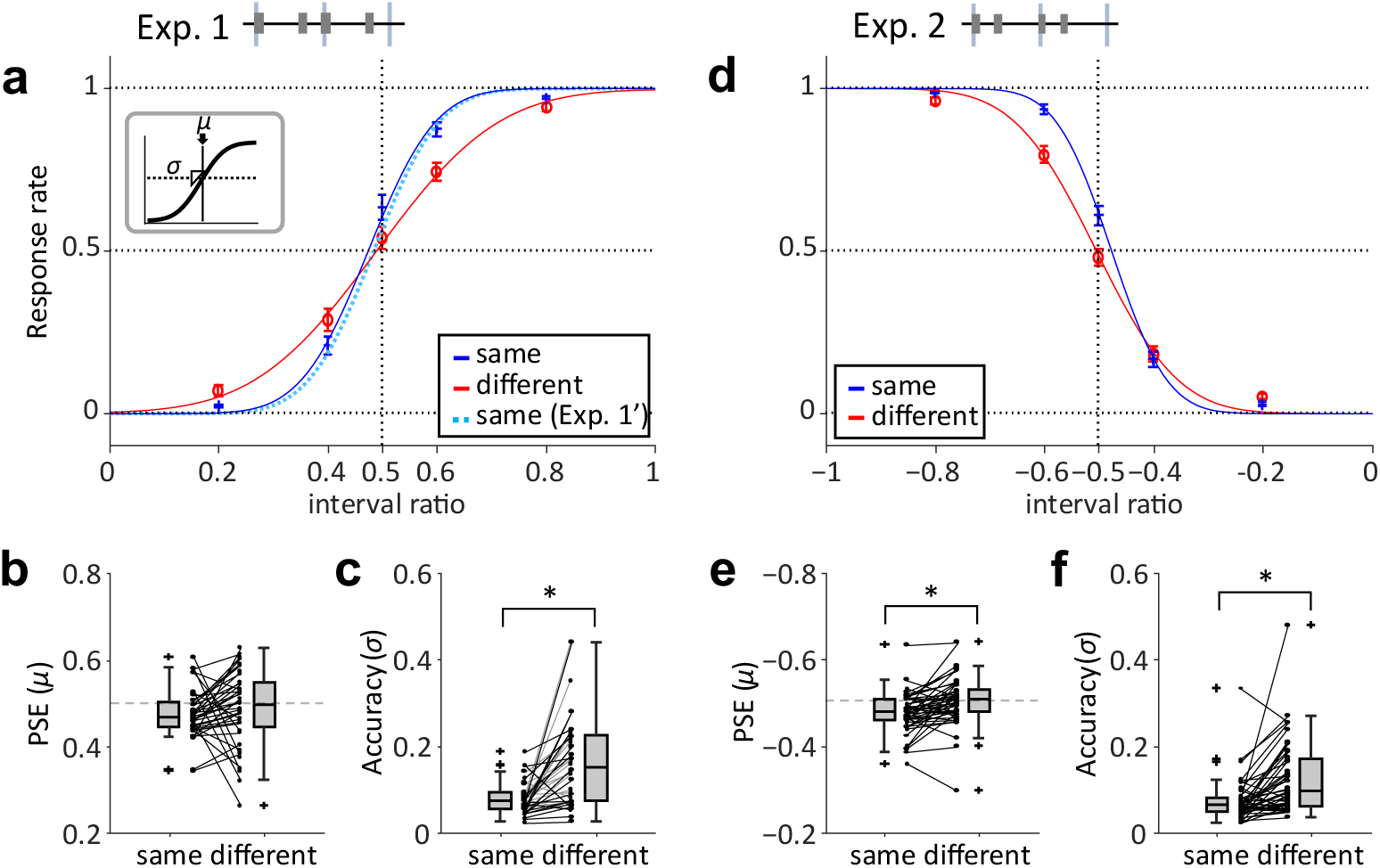
PSE and discrimination accuracy in long-short (Exp. 1 and 1’, a-c) and short-long rhythm (Exp. 2, d-f). (**a**) Response rates that comparison stimuli were “more bouncing” for each interval ratio. Red line indicates the different-frequency condition, and blue line indicates the same-frequency condition in Exp. 1. Light-blue dotted line indicates the same-frequency condition in Exp. 1’. Error bar shows standard error. Curves are sigmoidal approximations. (**b**) Distribution of PSE (*µ*) for same- and different-frequency conditions. (**c**) Distribution of discrimination accuracy (*σ*) for same- and different-frequency conditions. (**d**) Response rates that the comparison stimuli were “more bouncing” for each interval ratio. Red line indicates the different-frequency condition, and blue line indicates the same-frequency condition. Error bars are standard errors. Curves are sigmoidal approximations. (**e**) Distribution of PSE (*µ*) for same- and different-frequency conditions. (**f**) Distribution of discrimination accuracy (*σ*) for same- and different-frequency conditions. * indicates *p* < 0.05.

### Exp. 1&2: long-short rhythm (2:1), short-long rhythm (1:2)

We then assessed the effect of spectral differences in sound sequences on the discrimination accuracy of the long-short rhythm and short-long rhythm in Exp. 1 and 2, respectively. The accuracy was measured as the steepness (*σ*) of the fitted sigmoid curve. We screened the data according to the exclusion criteria and analyzed the remaining data obtained from 40 participants in Exp. 1 (**Figure 2a-c**). We calculated the response rate in each interval ratio and found that the steepness of fitted curve in the different-frequency condition was less than that in the same-frequency condition (**Figure 2a**). The estimated PSE (*μ*) (**Figure 2b**) was not significantly different between the two conditions (Wilcoxon signed-rank test, *Z* = 1.83, *p* = 0.068). In contrast, the discrimination accuracy (*σ*) (**Figure 2c**) was significantly larger in the different-frequency condition than in the same-frequency condition (*Z* = 4.73, *p* < 0.001).

We also analyzed the screened data obtained from 43 participants in Exp. 2 (**Figure 2d-f**), and found that the steepness in the different-frequency condition was less than that in the same-frequency condition (**Figure 2d**), similar to the long-short rhythm in Exp. 2. Both the estimated *μ* (**Figure 2e**) and *σ* (**Figure 2f**) were significantly greater in the different-frequency condition (*μ*: *Z* = 3.90, *p* < 0.001; *σ*: *Z* = 3.75, *p* < 0.001).

These results demonstrate that the discrimination accuracy decreases when the center frequency of narrow-band noises is alternated; thus, spectral inconsistency in the sound sequence deteriorates the discrimination of subdivided rhythms.

### Exp. 3: Straight Rhythm (1:1)

In Exp. 3, data from 30 participants survived after the screening. Different from Exp. 1 and 2, the response rate was assumed to be 0.5 at the regular interval (*r* = 0) and 0 at both ends (*r* = 1 and –1) because of the task design. We found again that the fitted curve in the different-frequency condition was less steep than that in the same-frequency condition (**Figure 3a**). The symmetric fitting showed that the estimated *σ* was significantly larger in the different-frequency condition (*Z* = 4.78, *p* < 0.001). The result indicates that the discrimination accuracy decreases with the spectral inconsistency even for the straight rhythm, similar to the long-short and short-long rhythms.

**Figure 3.**
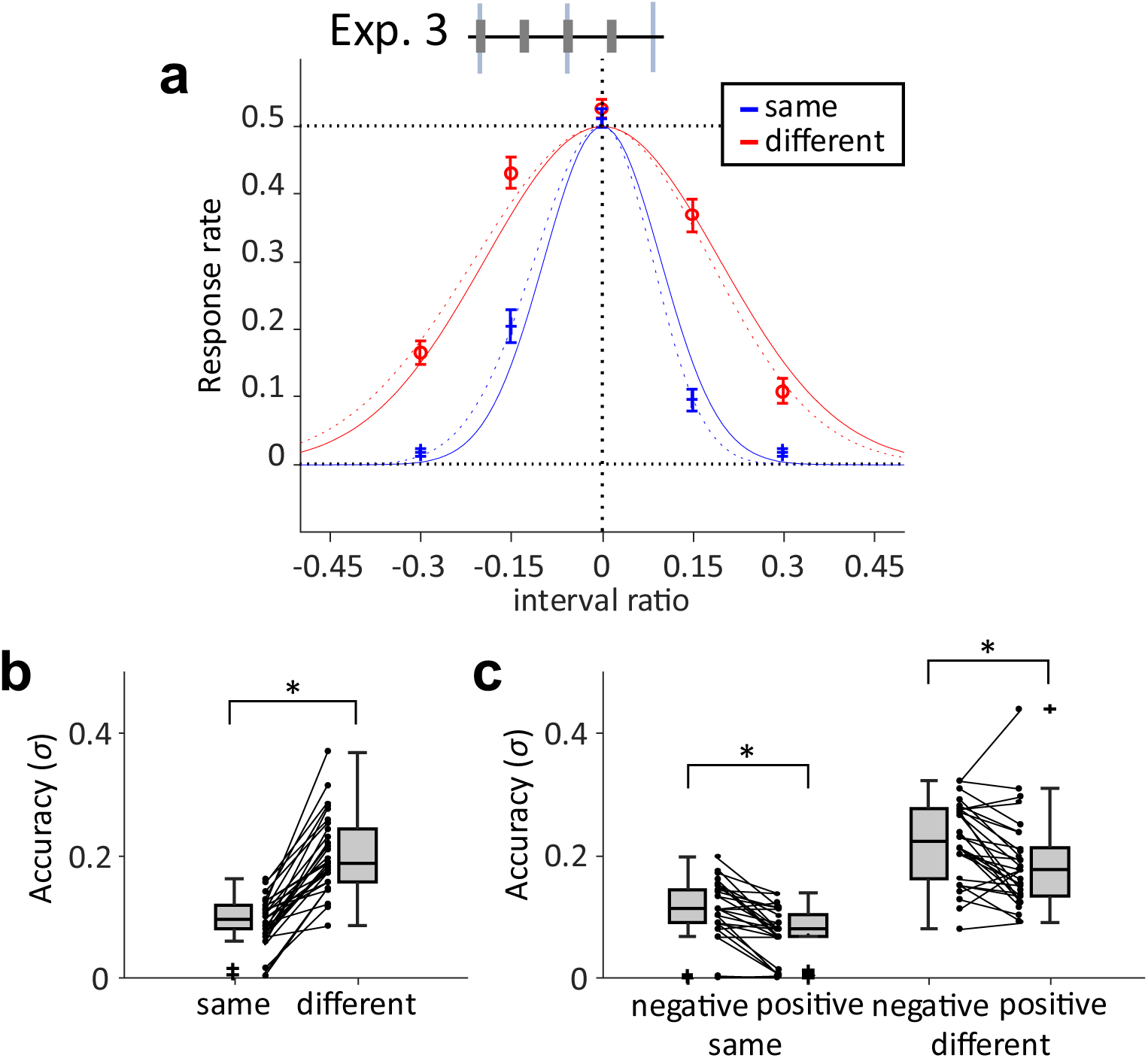
Discrimination accuracy in straight rhythm (Exp. 3). (**a**) Response rates that the comparison stimuli were “more bouncing” for each interval ratio. Red line indicates the different-frequency condition, and blue line indicates the same-frequency condition. Error bars are standard errors. Curves drawn by a solid line indicate symmetric fitting. Curves drawn by dotted line indicates asymmetric fitting. (**b**) Distribution of PSE (|*µ*|) for same- and different-frequency conditions. (**c**) Distribution of accuracy (*σ*) for same- and different-frequency conditions. * indicates *p* < 0.05.

We also performed the partial fitting for each of the positive and negative interval ratios to assess the asymmetry of the response rate data (shown as dotted lines in **Figure 3a**). The estimated *σ* of the asymmetric fitting (**Figure 3c**) was significantly larger for the negative interval ratio than for the positive ratio in both frequency: conditions (same-frequency: *Z* = 4.22, *p* < 0.001 different-frequency: *Z* = 3.18, *p* = 0.001).

### Effect of music experience

To examine the effect of musical instrument experience, participants in Exp. 1 were divided into two groups (with experiences: *n* = 23; without experiences: *n* = 20) by reporting whether they had instrumental experience, and σ was calculated for each frequency condition. The results showed a significant difference in σ between the frequency conditions in both groups (with experiences: *Z* = −3.47, *p* < 0.001; without experiences: *Z* = −3.54, *p* < 0.001). This suggests that discrimination accuracy decreased under the frequency conditions, regardless of musical experience.

## Discussion

The present study aimed to clarify how spectral consistency in sound sequences affects the perception of subdivision rhythm patterns. We assessed participants’ rhythm discrimination of band-pass noise sequences with manipulating sound timbre by differentiating their center frequencies. We found that discrimination accuracies (measured as *σ* of the fitted sigmoid function) decreased when the center frequencies alternated, irrespective of the rhythm pattern difference. This result indicates that the spectral consistency of sound sequences influences the accuracy of subdivided rhythm perception.

### Effect of spectral consistency on perceptual accuracy

Our analysis revealed that the discrimination accuracy decreased when the center frequency of the sequential band noises alternated (**Figure 2, 3**). This may be because a sound sequence that includes two spectral cues can be perceived as two segregated streams. Similarly, previous studies have reported that spectral differences promote stream segregation, which makes change detection more difficult for regular-interval (straight) rhythms [15,16], even though our results showed the same effect on non-straight rhythms. This decrease in perceptual accuracy can be quantified as the discrimination threshold. A study reported 6.2 ms displacement in the isochronous tone sequence with 130 ms intervals (corresponding to 260 ms IBI in our case) as a just-noticeable difference [10]. Another study estimated the threshold for discriminating swing from straight rhythms to be approximately 18 ms for 300-ms IBI [19]. We roughly estimated the thresholds in our data as interval ratios crossing 25% and 75% of the fitted curves. The mean estimated thresholds for the same-frequency condition in Exps. 1, 2, and 3 were 8.7, 7.1, and 9.6 ms, respectively. That for the different-frequency condition were 15.5, 10.7, and 19.5 ms, showing an apparent increase in the thresholds (corresponding to the decrease in perceptual accuracy) even though the estimated values were within the range of thresholds that were previously reported.

These findings suggest the following mechanism. Perceived timbre difference among sound elements produced by the center frequency manipulation enhances the segregation of auditory streams, and then the segregation elevates the threshold of discriminating subtle temporal shifts among sound elements across different streams, making the perceptual accuracy decrease. Note that we did not ask the participants to self-report whether they perceived sound streams as integrated or segregated. The precise relationship between our results and stream segregation should be addressed.

### Response asymmetry

The point of subjective equality (PSE) of the interval ratio in both Exps. 1 and 2 shifted towards 0 in the same-frequency condition (**Figure 2b, 2e**), indicating that participants perceived two stimuli equivalent when the comparison stimulus was slightly closer to the straight rhythm. The shift was not observed in the different-frequency condition. This effect may be due to the fixed presentation order of the standard and comparison stimuli in these experiments, e.g., short-term memory of the standard deformed before hearing the comparison; though such discussions were out of the scope of this study.

In Exp. 3, the estimated *σ* was significantly different between positive and negative interval ratios in both the same-frequency and different-frequency conditions (**Figure 3c**). This asymmetry can be explained by the time-shrinking effect [20]. When three consecutive tones with two onset intervals (T1 and T2), we tend to perceive the second time interval as shorter than the first (T2 < T1). This effect becomes weaker when the second interval is physically shorter than the first. A previous study showed that participants detected a negative shift of interval ratio more accurately than a positive shift in 1:2 rhythm [21], which is consistent with our result. Thus, when the standard rhythm pattern is straight rhythm, the perceptual accuracy may be higher in the long-short rhythm than in the short-long rhythm. However, it is also possible that a shift in PSE may have occurred during the experiment. Since the fitting was performed assuming PSE to be 0 in Exp. 3, further experiments are necessary.

### Individual variability

Our analysis did not find a significant effect of musical expertise. Contrary to our result, several studies have suggested that musical experience increases the accuracy of rhythm perception. For instance, people who have high musical experiences or music experts detect smaller changes in regular interval patterns than people with low musical experience [22, 23]. This discrepancy may be because no experts were recruited in our study.

One might wonder if participants exhibit consistent tendencies across different experiments. We started three online experiments at almost the same time, and some participants have joined two or three of them (see Participants subsection in Methods). Thus, we additionally performed *ad hoc* analysis for this issue by comparing individual discrimination abilities between Exp. 1 and 2 for people who participated in both experiments. We used less strict criteria for screening to increase the sample size (*n* = 21), in which we allowed to include participants who made a incorrect response once in the screening trials either in Exp. 1 or 2. As a result, almost all participants showed consistent tendency: both |*μ*| and *σ* were larger in the different-frequency condition than in the same-frequency condition (**Figure 4**), in both Exp. 1 (|*μ*|_different_−|*μ*|_same_: M ± SD = 0.032 ± 0.072; *σ*_different_−*σ*_same_: M ± SD = 0.040 ± 0.068) and Exp. 2 (|*μ*|_different_−|*μ*|_same_: M ± SD = 0.039 ± 0.056; *σ*_different_−*σ*_same_: M ± SD = 0.025 ± 0.075). These indicate consistent tendencies in PSE and discrimination accuracy across different experiments.

**Figure 4.**
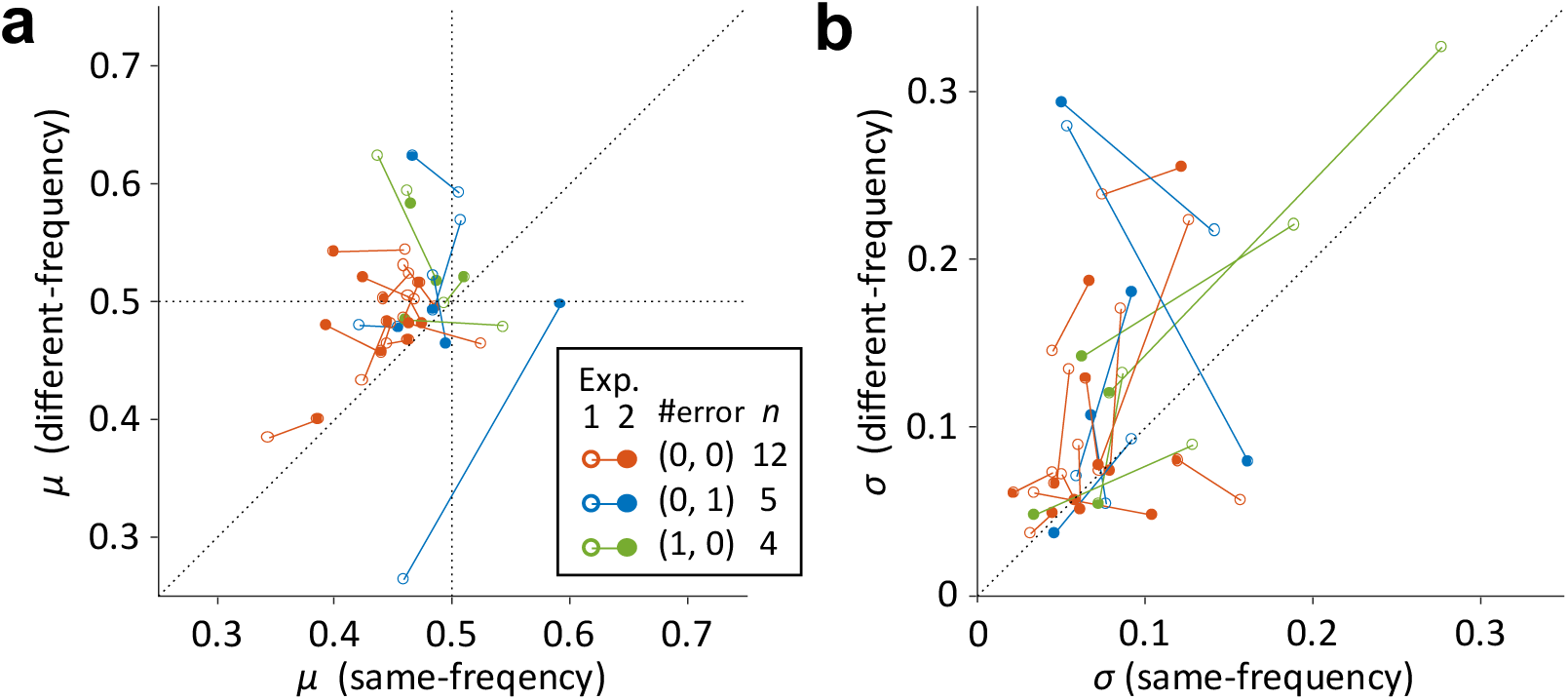
Comparison of *μ* and *σ* for overlap participants in Exp. 1 (a) and 2 (b). Scatterplots of the absolute value of PSE (|*μ*|) in the different-frequency condition for each |*μ*| in the same-frequency condition. Open circles indicate the values in Exp. 1, and filled circles indicate the values in Exp. 2. Red indicates all correct answers to the screening trials, blue indicates one incorrect answer to the screening trials in Exp. 1, and green indicates one incorrect answer to the screening trials in Exp. 2. Solid lines connect the same participants.

### Reliability of online experiment

The acoustic environment was variable among participants since they potentially heard stimuli from different audio devices with various background noises. We tried controlling such acoustic factors by introducing the headphone check and instructing them to join the experiment in a silent place. Further, we screened data according to the screening criteria, *i.e*., performance of screening trials and *R*^2^ of the sigmoid approximation. These criteria were determined arbitrarily, although we confirmed their validity as follows. We designed screening trials with clearly different interval ratios between standard and comparative stimuli to detect reliable participants who performed all these trials correctly. The *R*^2^ criteria were useful to remove data that exhibited extraordinary response profiles, e.g., a linear or U-shape pattern, and were far from the sigmoidal shape. We successfully removed such data at *R*^2^ = 0.5 with keeping a sufficient sample size. These criteria were rather conservative (we omitted 58.9% of data in total) but necessary to ensure the reliability of the data. The close match in the sigmoid approximation curves between Exps. 1 and 1’ (in-laboratory experiment) suggested further reliability of results obtained from our online experiments.

### Implications for cultural aspects of music

Our study also provides implications for the cultural aspects of music. The findings suggest that a decrease in perceptual accuracy may also occur in the musical performance since rhythmic patterns usually consist of multiple musical instruments. For example, drummers often use hi-hats and ride cymbals with the same center frequency in jazz music, where the swing ratio is considered an essential factor. The audience may feel more swing by composing rhythmic patterns with such instruments. However, many other parameters are involved in the actual rhythmic pattern, such as intensity, accents, and duration. For more accurate implications, it would be essential to investigate the effects of these factors on the accuracy of rhythm perception.

A previous study found a relationship between the perceptual accuracy of interval ratio and nationality [24]. For example, the accuracy of American participants was high only in the 2:1 ratio, while that of Turkish was high in 2:1 and 3:2, which were familiar swing ratios for them. This suggests perceptual accuracy may vary depending on culture-specific listening experiences and acquired musical knowledge. Therefore, it will be essential to investigate in future studies how the properties of rhythm perception, observed in the present study, vary with musical experience and belonging to culture to understand how we acquire rhythm perception.

Our paradigm can also be used to examine the effect of cultural differences on perceptual accuracy. For example, prior probability distributions over integer ratio on musical rhythm differ by cultural experience [25]. This may be due to the varying accuracy of discrimination depending on the cultural experience. A similar online experiment recruiting participants from different cultural backgrounds may provide mechanistic insights into the cultural evolution of music.

## Conclusion

We found that differences in center frequency in band noise lower the perceptual accuracy of subdivided rhythm regardless of the rhythmic pattern. This indicates that rhythm perception is affected not only by temporal structure but also by spectral parameters. Our findings bridge the gap between rhythm perception and auditory stream segmentation studies. Our paradigm also has the potential to examine the mechanism of cultural differences in music.

## Acknowledgment

This work was supported by JSPS KAKENHI Grant Number JP20K21804. We thank Dr. Kazutoshi Kudo for his helpful comments on early manuscripts. Furthermore, we want to express our gratitude to the colleagues from Okanoya Laboratory who provided insight and expertise that greatly assisted the research.

## Notes

### Competing Interest Statement

The authors have declared no competing interest.

### Summary of Updates

Figure 2 revised; Discussion on individual variability updated.

