## Supplementary material for "Spectral consistency in sound sequence affects perceptual accuracy in discriminating subdivided rhythmic patterns": Table S1

### Supporting information

***Supplementary analysis to assess the effect of high-low orders of the stimuli***

We used the linear mixed model as follows: $Y \sim frequency + order + frequency \times order + (1|participant)$. $Y$ indicates independent parameter, $\mu$ (PSE) or log $\sigma$ (discrimination accuracy), *frequency* indicates dummy variable representing center-frequency conditions (0: same-frequency; 1: different-frequency), *order* indicates dummy variable representing high-low order (0: high-low; 1: low-high), and $(1|participant)$ indicates random effect for each participant. We logarithmically transformed $\sigma$ to treat it as a parametric variable. After screening, participants were divided into two groups roughly in half (Exp. 1: $n_{\text{high-low}} = 19$, $n_{\text{low-high}} = 21$; Exp. 2: $n_{\text{high-low}} = 22$, $n_{\text{low-high}} = 21$; Exp. 3: $n_{\text{high-low}} = 12$, $n_{\text{low-high}} = 18$). We performed the analysis in R version 4.3.1. To fit the model, we used 'lmer' function of the lme4 and lmerTest package. We also adopted 'r.squaredGLMM' function of the MuMIn package to calculate the goodness of fit with R-squared value.

The analysis result showed that the effect of high-low order was not significant whereas $\sigma$ was significantly higher in the different-frequency condition (Table S1). This effect was consistent across three experiments. This suggests that only the center-frequency condition has an effect on discrimination accuracy. The model fitting performance was moderate (Exp.1: $R^2 = 0.52$; Exp.2: $R^2 = 0.51$; Exp. 3: $R^2 = 0.56$). These suggest that neither high-low order nor interaction effects contribute to discrimination accuracy.

Table S1. Summary of the linear-mixed model fitted to $\sigma$.

| **Experiment** | **Variable** | ***slope*** | ***SE*** | ***df*** | ***t*-value** | ***p*-value** |
|---|---|---|---|---|---|---|
| Exp. 1 | frequency | 0.26 | 0.06 | 38.0 | 4.33 | $p < 0.001$ |
| | order | 0.11 | 0.08 | 68.1 | 1.55 | $p = 0.126$ |
| | frequency×order | 0.02 | 0.09 | 38.0 | 0.19 | $p = 0.847$ |
| Exp. 2 | frequency | 0.14 | 0.05 | 41.0 | 2.51 | $p = 0.016$ |
| | order | 0.08 | 0.07 | 71.6 | 1.10 | $p = 0.277$ |
| | frequency×order | 0.12 | 0.08 | 41.0 | 1.49 | $p = 0.144$ |
| Exp. 3 | frequency | 0.34 | 0.07 | 28.0 | 4.77 | $p < 0.001$ |
| | order | −0.06 | 0.09 | 50.9 | −0.59 | $p = 0.558$ |
| | frequency×order | 0.11 | 0.11 | 28.0 | 0.97 | $p = 0.341$ |
